# Cryo-EM structure of monomeric CXCL12-bound CXCR4 in active state

**DOI:** 10.1101/2023.12.21.572776

**Authors:** Yezhou Liu, Aijun Liu, Xinyu Li, Qiwen Liao, Weijia Zhang, Lizhe Zhu, Richard D. Ye

## Abstract

CXC chemokine receptor 4 (CXCR4) and its chemokine ligand CXCL12 are crucial to embryonic development, bone marrow retention of hematopoietic progenitor cells, cancer metastasis, angiogenesis and HIV-1 infection. Yet the structural basis for CXCR4 recognition of full-length CXCL12 remains unknown despite available structures of CXCR4 in complex with small molecule antagonists and a viral chemokine. Here we present a cryo-EM structure of monomeric CXCL12-CXCR4 in complex with heterotrimeric Gi proteins at a resolution of 2.65 Å. The CXCL12-CXCR4 interaction at their N termini is stabilized by a polar interaction between the PC motif (C28^NT^ in CXCR4) and CXC motif (the unique P10 in CXCL12). The N terminal 8 amino acids in CXCL12 insert into the transmembrane binding pocket (chemokine recognition site 2, CRS2) of CXCR4 with S4 defining an upward turn of V3, P2 and K1. Polar interactions involving C186^ECL2^ and D187^ECL2^, along with hydrogen bonding with E288^7,39^ and D262^6,58^ stabilize the N terminus of CXCL12 in CRS2. Hydrophobic interactions between side chains aligning CRS2 and P2, L5 and S6 of CXCL12 further strengthen its binding to the receptor. CXCL12 inserts deep into the binding pocket, and the 3.2Å distance measured between V3 and the ‘toggle switch’ W^6.48^ for G protein activation is among the shortest of all chemokine-receptor pairs. Our findings provide structural insights into the recognition mechanism of CXCR4 for its chemokine ligand CXCL12.

## Introduction

Chemokines and their corresponding receptors are master regulators of immune functions under physiological and pathological conditions, including leukocyte migration, inflammation, immune surveillance, and wound healing (Griffith et al., 2014; Kufareva et al., 2017; Sokol & Luster, 2015; Viola & Luster, 2008).

Chemokines are small proteins (8 – 10 kDa) with a globular core and an extruding N-terminus, and can be classified into four subfamilies, CC, CXC CX_3_C and XC based on their conserved N-terminal cystine residues (Bachelerie et al., 2014; Hughes & Nibbs, 2018). All chemokine receptors belong to rhodopsin-like G protein-coupled receptors (GPCRs) with seven transmembrane (TM) helices and functional coupling to the Gi class of heterotrimeric G proteins for chemotaxis (Bachelerie et al., 2014; Hughes & Nibbs, 2018). With more than 50 chemokines and 20 receptors identified to date, the chemokine system constitutes a complex cell signaling network with significant promiscuity in ligand recognition (Anderson et al., 2016; Corbisier et al., 2015). An important aspect of such promiscuity could be attributed to the ligand-receptor interaction, which adopts a “two-site” mechanism involving multiple regions in the ligand as well as the receptor despite variability in the structures of chemokine and chemokine receptors (Ishimoto et al., 2023; Kufareva et al., 2017; Liu et al., 2020; Qin et al., 2015; Wu et al., 2010; Yen et al., 2022). Chemokine recognition site (CRS) 1 is defined as the receptor N terminus that interacts with the globular core of chemokines, and CRS2 is the transmembrane pocket that interacts with the N terminus of chemokines. CRS1 is crucial for its role in chemokine recruitment and association, whereas CRS2 is essential for receptor activation. With some chemokine receptors, a PC-motif in the receptor N terminus termed CRS1.5 is responsible for interaction with the N-loop-β3-strand disulfide bridge of the chemokine (Kleist et al., 2016; Siciliano et al., 1994; Zhang et al., 2021). The extensive receptor-chemokine interface allows multiple chemokines to promiscuously interact with the receptor extracellular loops while maintaining bound to the chemokine recognition sites (Yen et al., 2022). Such promiscuity allows multiple chemokines to interact with one receptor or another, constituting a complex regulatory network that ensures fine tuning of leukocyte functions.

CXCL12, originally identified as stromal cell-derived factor-1 (SDF-1), is the only endogenous chemokine ligand of CXCR4 (Bleul et al., 1996; Oberlin et al., 1996). The CXCL12/CXCR4 axis plays important physiological functions. In early development, CXCR4 acts as a master switch of embryonic development where progenitor cells migrate to their destinations then differentiate into tissues and organs. Throughout mammalian lifespan, CXCR4 is responsible for lymphocyte and leukocyte trafficking (De Filippo & Rankin, 2018), homeostasis of bone marrow immune cell niche (Sugiyama et al., 2006), infection and inflammation (Doranz et al., 1999; Feng et al., 1996; Salvatore et al., 2010) as well as cancer metastasis (Burger & Peled, 2009; Drury et al., 2011; Yang et al., 2007). CXCR4 also acts as a co-receptor of HIV-1 infection (Doranz et al., 1999; Feng et al., 1996).

These findings indicate that CXCR4 is responsible for the majority of biological functions of CXCL12, although this chemokine also bind to the atypical chemokine receptor ACKR3 (CXCR7) that mediates β-arrestin signaling and constitutive internalization of CXCL12 (Decaillot et al., 2011; Huynh et al., 2020; Werner et al., 2011). Given the important role of CXCR4 in various physiological and pathological conditions, effort has been made in understanding the structural basis of CXCL12 and CXCR4 interactions. To date, three CXCR4 structures have been available (Qin et al., 2015; Wu et al., 2010), and a cryo-EM structure of CXCL12-ACKR3 has also been resolved (Yen et al., 2022). Wu *et al* first reported two crystal structures of CXCR4 in complex with a small molecule antagonist IT1t and a cyclic peptide antagonist CVX15, respectively (Wu et al., 2010). The CXCR4 receptor homodimers were found in these structures, in a 2:1 stoichiometry to the ligands. Handel and colleagues solved the crystal structure of CXCR4 in a 1:1 complex with the viral chemokine vCCL2 (vMIP-II) (Qin et al., 2015). vCCL2 is derived from herpesvirus HHV-8 (Kaposi’s sarcoma-associated herpesvirus) and functions as a broad-spectrum antagonist for several chemokine receptors (Kledal et al., 1997). vCCL2 is a high affinity antagonist for CXCR4 (Zhou et al., 2000). As a result, these three structures show CXCR4 in an inactive conformation. More recently, Handel and coworkers (Yen et al., 2022) reported a cryo-EM structure of ACKR3-CXCL12 that displayed a distinctive binding mode. In their model, CXCL12 is rotated by 80° and shifted by ∼10 Å towards TM5 and TM6, thus altering its interaction with CRS1 and CRS1.5 of ACKR3. The CRS2 recognition is mostly retained but signaling through ACKR3 is biased towards the β-arrestin pathway (Yen et al., 2022). In addition to these structures, several computational models of CXCL12-CXCR4 interaction were proposed (Gustavsson et al., 2019; Kufareva et al., 2014; Ngo et al., 2020; Stephens et al., 2020; Wescott et al., 2016). However, the discrepancies among the computational models and mutational studies call for an experimentally solved CXCR4-CXCL12 structure in activated conformation.

Here, we present a cryo-EM structure of CXCR4-CXCL12 complex bound to heterotrimeric Gi proteins at an overall resolution of 2.65 Å. Analysis of the structure identified key amino acid residues at the CXCR4 chemokine recognition sites for CXCL12 binding, receptor activation and transmembrane signaling. By combining functional assays with site-directed mutagenesis of CXCR4 and molecular dynamics (MD) simulations, we further verified the interactions between CXCR4 and CXCL12. Polar interactions characterize the binding between the chemokine in its N terminus and CRS2 of CXCR4. A hydrophobic interaction network is observed, extruding from CRS1 to the binding pocket at CRS2 and the intracellular loops, for receptor conformational changes towards an active state for G protein activation and downstream signaling. Altogether, these data provide new insights into the CXCR4-CXCL12 interaction and a structural basis for CXCR4 activation, that may aide in future design of novel ligands targeting the CXCL12/CXCR4 axis.

## Methods

### Constructs

The coding sequence of wild-type human CXCR4 was synthesized by General Biol (Chuzhou, China) and was cloned into the pFastBac transfer plasmid (Invitrogen, Carlsbad, CA). Heat-stabilized BRIL, FLAG tag and NanoBiT tethering method was used to facilitate expression and purification (Chun et al., 2012; Duan et al., 2020). This strategy is widely used in enhancing GPCR recombinant expression and generally does not affect receptor activity (Chen et al., 2022; Lv et al., 2016; Zou et al., 2012). All constructs were generated using Phanta Max Super-Fidelity DNA polymerase (Vazyme Biotech, Shanghai, China; Cat# C115) and verified by DNA sequencing (Genewiz, Suzhou, China). A dominant-negative (DN) Gαi format containing G203A/A326S mutations was engineered to increase the stability of the Gαiβ1γ2 complex (Liu et al., 2016). The coding sequence of CXCL12 and scFv16 fused with a GP67 signal peptide at the N-terminus and an 8×His tag at the C terminus were cloned into the pFastBac vector. For G protein dissociation assay and cAMP assay, the coding sequences of single-point mutants were synthesized by General Biol.

### Insect cell expression

Baculoviruses were prepared using the Bac-to-Bac Expression System (Invitrogen) and expressed using the baculovirus method. *Spodoptera frugiperda* Sf9 insect cells (Invitrogen) were grown in suspension at 27°C, when the cell density reached 2×10^6^ cells per ml, human CXCR4, CXCL12, Gαi, Gβ1, Gγ2 were coinfected at a ratio of 1:2:1:1:1. After 48 h of infection, cells were collected by centrifugation and then stored at −80°C until use. The baculovirus of scFv16 was prepared in the same way as for CXCR4. *Trichoplusia ni* Hi5 insect cells were grown to a density of 2.5 × 10^6^ per ml and infected with virus at a ratio of 1:40. After 60 h, the supernatant was collected for purification.

### Purification of ScFv16

Secreted scFv16 was purified from expression media of baculovirus-infected Sf9 insect cell culture using Ni-NTA. The supernatant from 2 L of culture was collected and loaded onto a gravity column of Ni-NTA resin. The resin was washed with 20 mM HEPES pH 7.5, 500 mM NaCl, and the protein was eluted by 20 mM HEPES pH 7.5, 150 mM NaCl and 250 mM imidazole. Eluted protein was concentrated and loaded onto Superdex 200 increase 10/300 size exclusion column (GE Healthcare Life Sciences, Sweden). The peak fractions were collected and concentrated, fast-frozen in liquid nitrogen, and stored at −80 °C.

### Complex purification

Cell pellets from 1 L of culture were thawed at room temperature and suspended in the buffer containing 20 mM HEPES pH 7.5, 50 mM NaCl, 5 mM CaCl_2_, 5 mM MgCl_2_, 25 mU ml^−1^ apyrase (Sigma, Cat. No. A6535), 10 μM CXCL12 with 100 × concentrated EDTA-free protease inhibitor cocktail (TargetMol). Subsequently, 0.5% (w/v) n-dodecyl-β-d-maltoside (DDM, Anatrace) and 0.1% (w/v) cholesteryl hemisuccinate (CHS, Anatrace) were added to solubilize complexes for 2 h at 4 °C. Insoluble material was removed by centrifugation at 30,000 × g for 30 min and the supernatant was incubated with anti-FLAG affinity resin (GenScript Biotech, Piscataway, NJ). After that, the resin was loaded onto a gravity-flow column and washed with 20 column volumes of 20 mM HEPES pH 7.5, 0.1% DDM, 0.01% CHS, 100 mM NaCl, 2 mM CaCl_2_, 1 μM CXCL12. Finally, the complex was collected in buffer containing 0.2 mg/ml FLAG peptide and concentrated to 400 μL in an Amicon® Ultra-15 Centrifugal Filter Unit (Millipore, Burlington, MA). Complexes were loaded onto a Superose 6 Increase 10/300GL column (GE Healthcare) followed by Superdex 200 Increase 10/300 column (GE Healthcare) with buffer containing 20 mM HEPES pH 7.4, 100 mM NaCl, 0.00075% (w/v) LMNG, 0.0002% (w/v) CHS to separate complex from contaminants. Eluted fractions consisting of CXCR4-Gi complex were pooled and concentrated before being flash frozen in liquid nitrogen and stored at −80°C.

### Cryo-EM grid preparation and data collection

For cryo-EM grids preparation, 3 μL of the purified complex were loaded onto a glow-discharged holey grid (Quantifoil Au 300 mesh R1.2/1.3), and subsequently were plunge-frozen in liquid ethane using Vitrobot Mark IV (Thermo Fisher Scientific, Waltham, MA). Cryo-EM imaging was collected at the Kobilka Cryo-EM Center of The Chinese University of Hong Kong, Shenzhen, on a Titan Krios at 300 kV using Gatan K3 Summit detector with a pixel size of 0.85 Å. Inelastically scattered electrons were excluded by a GIF Quantum energy filter (Gatan, USA) using a slit width of 20 eV. Images were taken at defocus ranging from −1.0 to −2.5 μm using the semi-automatic data acquisition software SerialEM. A total of 12,322 image stacks were collected in 96 hours with total dose of 52 e/Å^2^ over 2.5 s exposure on each movie.

### Cryo-EM image processing and map construction

The single particle analysis of CXCR4-Gαi complexes was performed with cryoSPARC v3.3.1 (Structura Biotechnology Inc., Toronto, Canada). The image stacks were firstly subjected to patch motion correction and patch CTF estimation. 3,370,020 particles were auto-picked and then subjected to 2D classification and *ab initio* reconstruction. After multiple rounds of hetero-refinement, a final set of 523,651 particles were subject to non-uniform refinement and local refinement, yielding a map with an overall resolution of 2.65 Å at a Fourier shell correlation of 0.143.

### Model building and refinement

Predicted model of active-state CXCR4 receptor from AlphaFold2 were used as initial model for rebuilding and refinement against the electron microscopy density map. UCSF Chimera-1.1470 was used to dock the model into the electron microscopy density map, and followed by iterative manual adjustment and rebuilding in COOT-0.9.8. Then models were further refined and validated in Phenix-1.2073 programs (Supplementary Table 1). Structural figures were generated using UCSF Chimera-1.14, ChimeraX-1.274 and PyMOL-2.0.

### Mutagenesis study

CXCR4 cDNA in the pcDNA3.1(+) vector (Invitrogen) was used as a template for point mutation. The mutations were introduced into the coding sequence of the receptor at designated sites by chemical synthesis and overlap extension PCR (General Biol), then cloned into pcDNA3.1(+) vectors. All constructs were verified by sequencing (Genewiz) and transfected into cells for transient expression using Lipofectamine 3000 (Invitrogen).

### cAMP inhibition assay

For cAMP inhibition assay, wild-type CXCR4 and its mutants were transiently expressed in HeLa cells 24 hrs prior to experiment. Cells were resuspended in HBSS buffer with 5 mM HEPES, 0.1% BSA (w/v) and 0.5 mM 3-isobutyl-1-methylxanthine (IBMX). Different concentrations of CXCL12 (#30028A; Peprotech, Rocky Hill, NJ) were prepared in the abovementioned stimulation buffer. Cells were treated with 2.5 μM forskolin and CXCL12 at different concentrations for 30 mins in a cell incubator. Intracellular cAMP level was measured using LANCE Ultra cAMP kit (#TRF0263; PerkinElmer Life Sciences, Waltham, MA) following the manufacturer’s instructions. Signals of time resolved-fluorescence resonance energy transfer (TR-FRET) were collected by an EnVision 2105 multimode plate reader (PerkinElmer). Intracellular cAMP levels were calculated as instructed by the manufacturer.

### NanoBiT-based G protein dissociation assay

Dissociation of G protein heterotrimer was examined with a NanoBiT-based G protein dissociation assay (Inoue et al., 2019). HEK293T cells were plated in a 24-well plate, then transfected with a mixture of 100 ng pcDNA3.1 vector encoding CXCR4 (WT/mutants), 100 ng pcDNA3.1 vector encoding Gαi1-LgBiT, 200 ng pcDNA3.1 vector encoding Gβ1 and 200 ng pcDNA3.1 vector encoding SmBiT-Gγ2 (per well in a 24-well plate), respectively. Twenty-four hours after transfection, cells were collected and resuspended in 20 mM HEPES HBSS. Cells were loaded onto a 384-well white plate at a volume of 20 μL and incubated with 10 μM coelenterazine H for 2 hrs at room temperature (Yeasen Biotech, Shanghai, China). Baseline chemiluminescence signals were read using an Envision 2105 multimode plate reader (PerkinElmer). CXCL12 at different concentrations were then added. Signals were measured after 15 mins’ incubation and divided by the baseline readout. The percentage changes of signals were further normalized by HBSS-treated signal and plotted as a function of different concentrations of CXCL12 based on three independent experiments, each with triplicate measurements.

### Surface expression analysis

HEK293T cells were transfected with WT or mutant CXCR4 expression plasmids for 24 hrs at 37°C. Then cells were harvested and washed in HBSS containing 5% BSA for three times on ice. The cells were then incubated with an APC-labeled anti-CXCR4 antibody (12G5; Thermo Fisher Scientific, Cat #17-9999-42; 1:50 diluted by HBSS buffer) for 30 mins on ice and washed with HBSS. The fluorescence signals demonstrating the antibody-receptor complex on the cell surface were quantified by flow cytometry (CytoFLEX, Beckman Coulter, Brea, CA).

### MD simulations

The atomic coordinates of CXCR4-CXCL12 complex were extracted from the cryo-EM structure solved in this study. Protonation state of the complex was assigned by the web server H++ assuming pH 7.4. Then the processed complex was embedded in a bilayer composed of 1-palmitoyl-2-oleoyl-sn-glycero-3-phosphocholine (POPC) lipids using CHARMM-GUI membrane builder (Lee et al., 2019). The system was solvated in a periodic 0.1 M NaCl TIP3P water box. The force fields of Amber ff19SB and lipid17 were used to model protein and membrane lipid atoms, respectively. The simulations were then performed on GROMACS (version 2021.3) using the GPU implementation. The system was energy minimized for 10,000 steps, then 200 ns of restrained MD simulation was performed to fully relax and equilibrate the system contained solvent, membrane, and the complex at 303.1 K, 1.0 bar using the particle mesh Ewald (PME) method, three independent 600-ns long production MD simulations was carried out. After production, three 600-ns MD trajectories were produced. The stability of the peptide-receptor complex was monitored by root mean square deviations (RMSD). The trajectories from three independent simulation analyses indicate that CXCL12 displayed a stable pose of hook shape in the orthosteric pocket of CXCR4.

### Statistical Analysis

The data were analyzed with statistical software Prism 9.5.0 (GraphPad, San Diego, CA). The dose-response curves were plotted as the log[agonist] vs. response equation (three parameters) in the software. For cAMP accumulation and G protein dissociation assays, data points were presented as mean ± SEM of the percentages of maximal responses for each sample from at least three independent experiments. The EC_50_ values were calculated from dose-response curves. For cell surface expression, data points were presented as the percentages (mean ± SEM) of the flow cytometry fluorescence signals of WT CXCR4. Analysis of Variance (ANOVA) using the one-way method was applied for statistical comparison. A *p* value of 0.05 or lower is considered statistically significant.

## Results and Discussion

### Molecular interactions between CXCR4 and CXCL12

Human CXCR4, CXCL12 (SDF-1α) and heterotrimeric Gi proteins were co-expressed in *Sf9* cells for complex formation as described in *Methods*. The complex of CXCR4, CXCL12 and Gi proteins were further stabilized by a single-chain variable antibody fragment scFv16 for specimen vitrification and single particle cryo-EM data collection (Fig. S1). The electron density map from cryo-EM data yielded a three-dimensional (3D) structural model of CXCR4-CXCL12-Gi complex at a global resolution of 2.65 Å (Fig. 1 A-C, Fig. S2). The electron density for CXCR4 is clearly defined in the seven transmembrane (TM) domains as well as the intra– and extra-cellular loops except that the N terminus of CXCR4 (a.a.1-24) was difficult to capture due to intrinsic instability (Fig. 1D, 1E).

**Figure 1.**
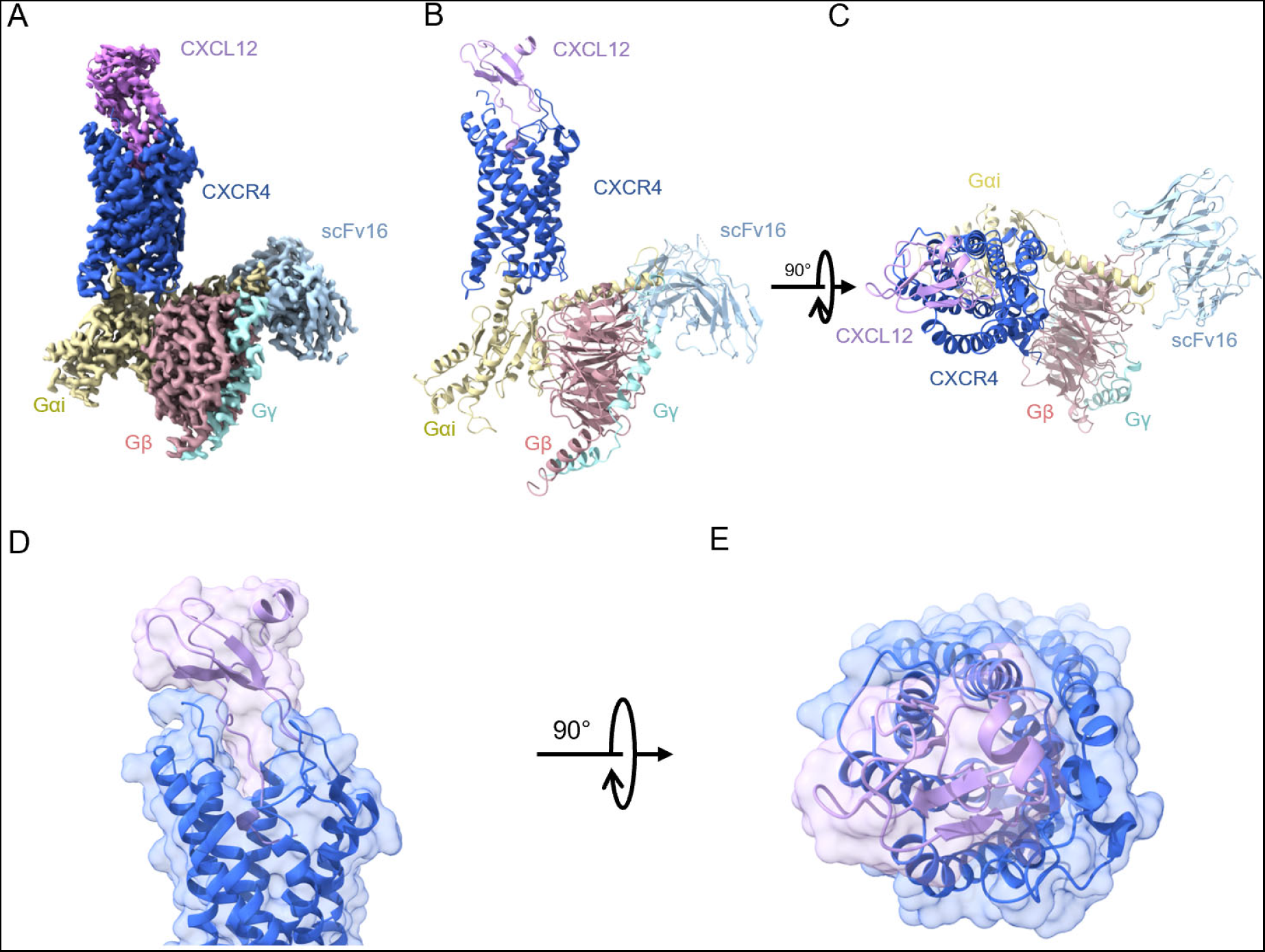
Overall structure of CXCR4-CXCL12-Gi complex. **A** Cryo-EM electron density map of CXCR4-CXCL12-Gi complex. CXCR4 is shown in blue, CXCL12 in lightpink, Gαi in yellow, Gβ in salmon, Gγ in cyan and scFv16 in slate blue. **B** Side view of the structure of CXCR4-CXCL12-Gi complex. **C** Extracellular view of the structure of CXCR4-CXCL12-Gi complex. **D** Interface between CXCR4 and CXCL12 (side view). Surface view is displayed transparently. **E** Interface between CXCR4 and CXCL12 (top view).

The tertiary structure of CXCL12 in this model covers a.a. 22-83 (without the N terminal signal peptide of 21 a.a. and the C terminal 6 a.a.) as shown in Fig. 2A, with different parts of the CXCL12 primary sequence colored for clarity. The folding of CXCL12 is consistent with the NMR structure of CXCL12 monomer (Veldkamp et al., 2009; Yen et al., 2022; Ziarek et al., 2017). The various parts of CXCL12 include an N terminus loop (grass green), a CXC motif (yellow) followed by an N-loop (orange), three β strands (β_1_, β_2_ and β_3_ strand) connected by a 30s-loop (light blue) and a 40s-loop (bright green) (Fig. 2A). Two pairs of disulfide bonds are present in CXCL12, between residue pairs C30-C55 and C32-C71. In line with previous discoveries towards the “two-site” mode of chemokine binding (Kleist et al., 2016; Scholten et al., 2012; Siciliano et al., 1994), the chemokine ligand *CXCL12* in our model interacts through its *globular core* with the N terminus of CXCR4 (CRS1), *CXC motif* with the PC motif of CXCR4 (CRS1.5), and *N terminus* with the TM binding pocket of CXCR4 (CRS2) (Fig. 2B). Due to intrinsic flexibility of the receptor N-terminus, the structure of CRS1 of CXCR4 is just resolved to K25^NT^. At CRS1.5, an intermediate region between CRS1 and CRS2, a prominent feature in this model is a polar interaction between the peptide backbone of C28^NT^ in CXCR4 and P10 of CXCL12, where proline defines the intervening “X” in the CXC motif (Fig. 2C). Proline in this position of CXCL12 is unique among all CXC chemokines. As discussed below, this interaction may help to orient the chemokine’s N terminus in its interaction with CRS2. The interface between CXCR4 and CXCL12 at CRS2 is defined by polar and nonpolar interactions. The chemokine N terminus inserts deeply into the receptor TM pocket with multiple polar contacts (Fig. 2D). The first four residues of CXCL12 take a horizontal pose in the binding pocket (Fig. 2D). K1 of CXCL12 forms polar interactions with C186^ECL2^ and D187^ECL2^ in extracellular loop 2 (ECL2) of CXCR4, and V3 of CXCL12 forms a hydrogen bond with E288^7.39^. S4 of CXCL12 has multiple polar interactions with H281^7.32^ and E288^7.39^, defining an upward turn from the horizontal pose towards the extracellular region. The remaining N terminus from L5 to R8 take a vertical pose in CRS2. In this segment, Y7 of CXCL12 has polar interactions with D187^ECL2^ of CXCR4, while R8 of CXCL12 forms a hydrogen bond with D262^6.58^ of CXCR4. The ligand interaction with CRS2 is also supported by a network of hydrophobic interactions between CXCL12 and several CXCR4 residues, including L41^1.35^, W94^2.60^, H113^3.29^, Y116^3.32^, I185^ECL2^, F189^ECL2^, Y255^6.51^, I259^6.55^, L266^6.62^, E268^6.64^, and S285^7.36^ (Fig. 2E). Specifically, P2 of CXCL12 has close hydrophobic interactions with W94^2.60^ and Y116^3.32^, L5 forms nonpolar interactions with adjacent L41^1.35^ and H113^3.29^, and S6 has short-range hydrophobic interaction with I185^ECL2^. These critical residues for agonistic chemokine binding identified at receptor CRS2 are mostly consistent with the first hypothetic model of CXCR4-CXCL12 interaction by molecular modelling (Qin et al., 2015). However, the majority of interaction details between the receptor and chemokine residues are different, especially the first four residues of CXCL12 which favor a position pointing towards TM2 in Qin’s model.

**Figure 2.**
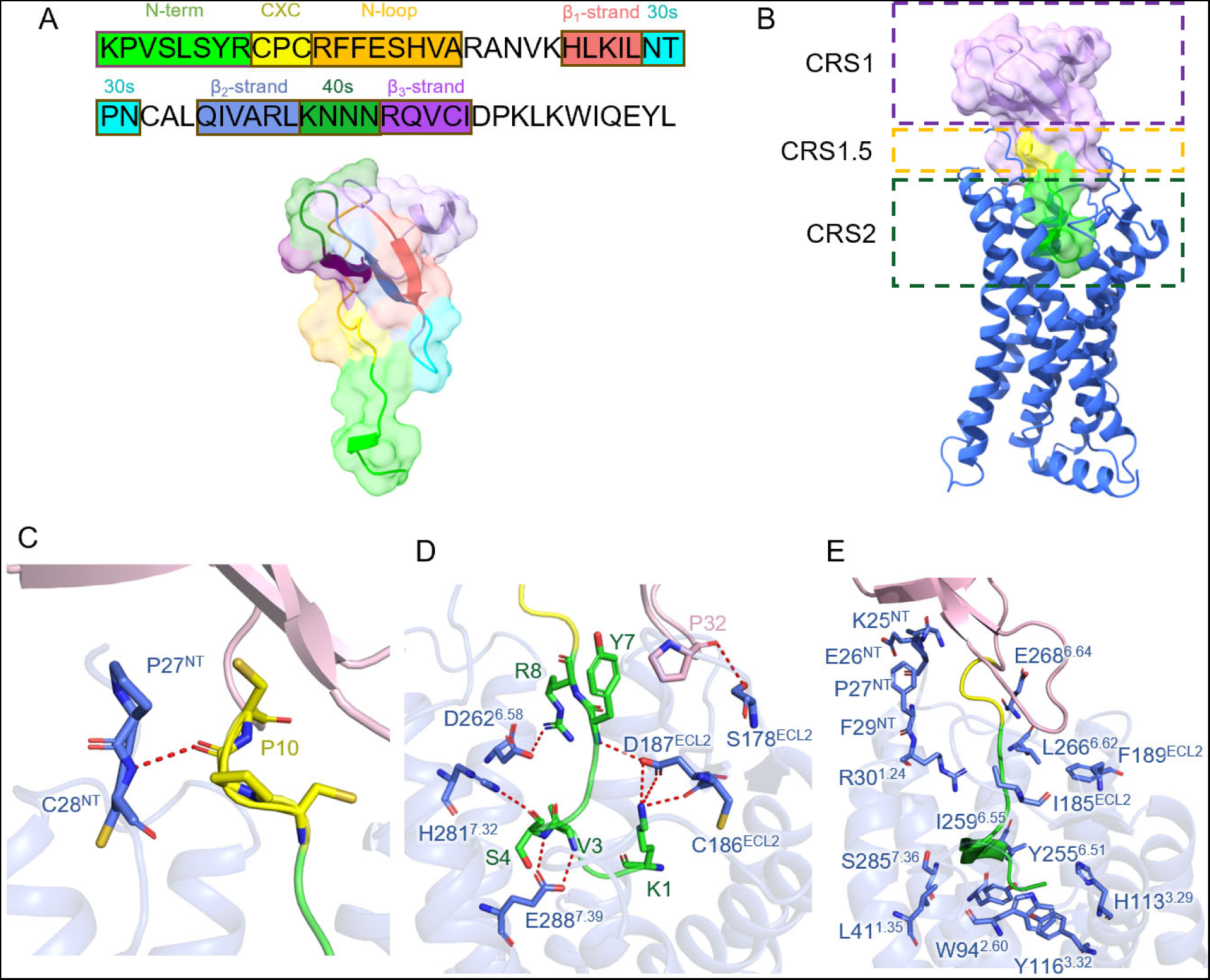
Structural details at chemokine recognition sites. **A** The sequence (upper panel) and the structure (lower panel) of CXCL12. N-term (light green): chemokine N-terminal K1-R8; CXC (yellow): CXC motif; N-loop(orange): N-terminal loop of CXCL12; β_1_-strand (salmon): the first β strand of CXCL12; 30s (cyan): 30s-loop connecting β_1_-strand and β_2_-strand; β_2_-strand (marine blue): the second β strand of CXCL12; 40s (grass green): 40s-loop connecting β_2_-strand and β_3_-strand; β_3_-strand (purple): the third β strand of CXCL12. **B** Side view of the structure of CXCR4-CXCL12 complex. The receptor is shown in blue cartoon, and the chemokine is displayed with cartoon and transparent surface. Green: N-term, yellow: CXC motif, lightpink: the globular core of CXCL12. Chemokine recognition sites CRS1, CRS1.5 and CRS2 are highlighted. **C** Interactions between CXCR4 and CXCL12 at CRS1.5. Polar interactions are highlighted by yellow dashes. **D** Polar interactions between CXCR4 and CXCL12 at CRS2, highlighted by yellow dashes. **E** Hydrophobic and other nonpolar interactions between CXCR4 and CXCL12.

In the extracellular region for chemokine recruitment, P32 is an important residue of CXCL12 in the 30s-loop connecting β1 and β2 strand. It interacts with S178^ECL2^ of CXCR4 with polar bonds, defining the chemokine recognition site 3 (CRS3) (Kleist et al., 2016). This P32-S178^ECL2^ interaction at CRS3 is consistent with previously reported computational models as well as cryo-EM models of other chemokine receptors (Lu et al., 2022; Shao et al., 2022; Urvas & Kellenberger, 2023), thereby supporting the stabilization of the ECL2 conformation. Multiple non-polar interactions at the receptor’s N terminus further enhance the stability of interaction between the globular core of chemokine and several residues of CXCR4, including K25^NT^, E26^NT^, P27^NT^ and F29^NT^ as well as I185^ECL2^ and F189^ECL2^. The hydrophobic interactions extending from the extracellular chemokine recognition sites to the TM domain of the receptor form a structural network to relay conformational changes upon chemokine binding (Wescott et al., 2016).

To further verify the structural information for CXCR4-CXCL12 interaction, we introduced site-directed mutagenesis at key residues for chemokine recognition. By applying NanoLuc Binary Technology (NanoBiT), we tested the CXCR4 mutants in functional assays for G protein dissociation (Fig.3 A-C) (Inoue et al., 2019), as well as inhibition of cAMP accumulation (Fig.3 D-F) that reflects Gi protein signaling in response to CXCL12 binding. The C28A mutant reduced the potency of CXCL12 by 2 orders of magnitude, suggesting the importance of the interaction between CRS1.5 of the receptor and the proline in the CXC motif that interacts with C28^NT^ for CXCR4 activation. For residues in ECL2, Ala substitution at C186^ECL2^ and D187^ECL2^ markedly reduced the functions of the receptor while Ala substitution of S178^ECL2^ (S178A) did not markedly affect G protein dissociation and cAMP accumulation (Fig. 3 B and E). Notably, Ala substitution at C186 significantly reduced the surface expression of CXCR4, whereas the expression level of D187A remains normal (Fig. S3). C186^ECL2^ is known to form disulfide bond with C109^3.25^ (Wu et al., 2010), a structural feature that stabilizes the global conformation of most Class A GPCRs. Another disulfide link is formed between C28^NT^ and C274^7.25^, which is a conserved feature among chemokine receptors except CXCR5 and CXCR6 (Kleist et al., 2016; Kufareva et al., 2017). These two disulfide bonds at the extracellular side of CXCR4 may be crucial for ligand binding by restricting the entry to CRS2. Deeper into the binding pocket, Ala substitutions at D262^6.58^, H281^7.32^ and E288^7.39^ significantly ameliorated the functions of CXCR4, as demonstrated by reduced G protein dissociation and completely abolished cAMP accumulation (Fig.3 C and F). The results from these functional assays support our structural model and together suggest that CXCR4 interaction with the globular core of CXCL12 may not be a determinant for receptor activation, while interactions of the N terminus of CXCL12 at CRS2 play crucial roles for CXCR4 activation.

**Figure 3.**
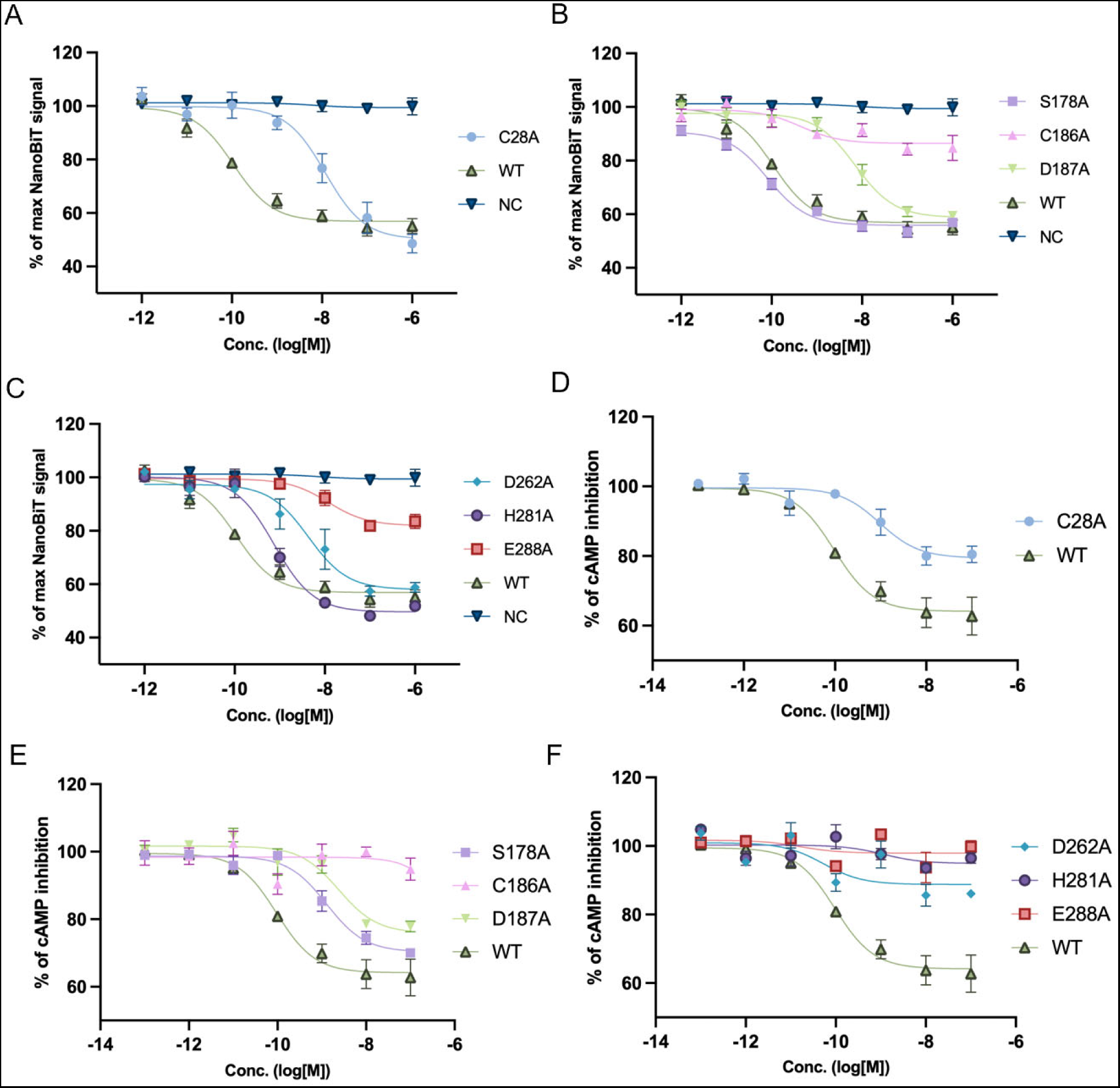
Functional examination of key residues at chemokine recognition sites. Alanine substitution was introduced for chemokine-interacting residues of CXCR4. NanoBiT-based G protein dissociation assay was performed for mutants lining **A** CRS1.5 and **B-C** CRS2. cAMP inhibition assay was conducted for mutants in **D** CRS1.5 and **E-F** CRS2. WT: wild-type CXCR4. NC: empty vector control. Data were collected from three independent experiments, each with three replicates.

### Thermodynamic features at key binding residues between CXCR4 and CXCL12

To further evaluate the interactions between CXCR4 and CXCL12, we performed full-atom molecular dynamics (MD) simulations of the complex at room temperature. The cryo-EM structure of the CXCR4-CXCL12 complex was overall stable under thermodynamic perturbation. The trajectories from three independent simulations analyses show that CXCL12 displayed a stable pose of hook shape in the orthosteric pocket of CXCR4 (Fig. 4A, Fig. S4).

**Figure 4.**
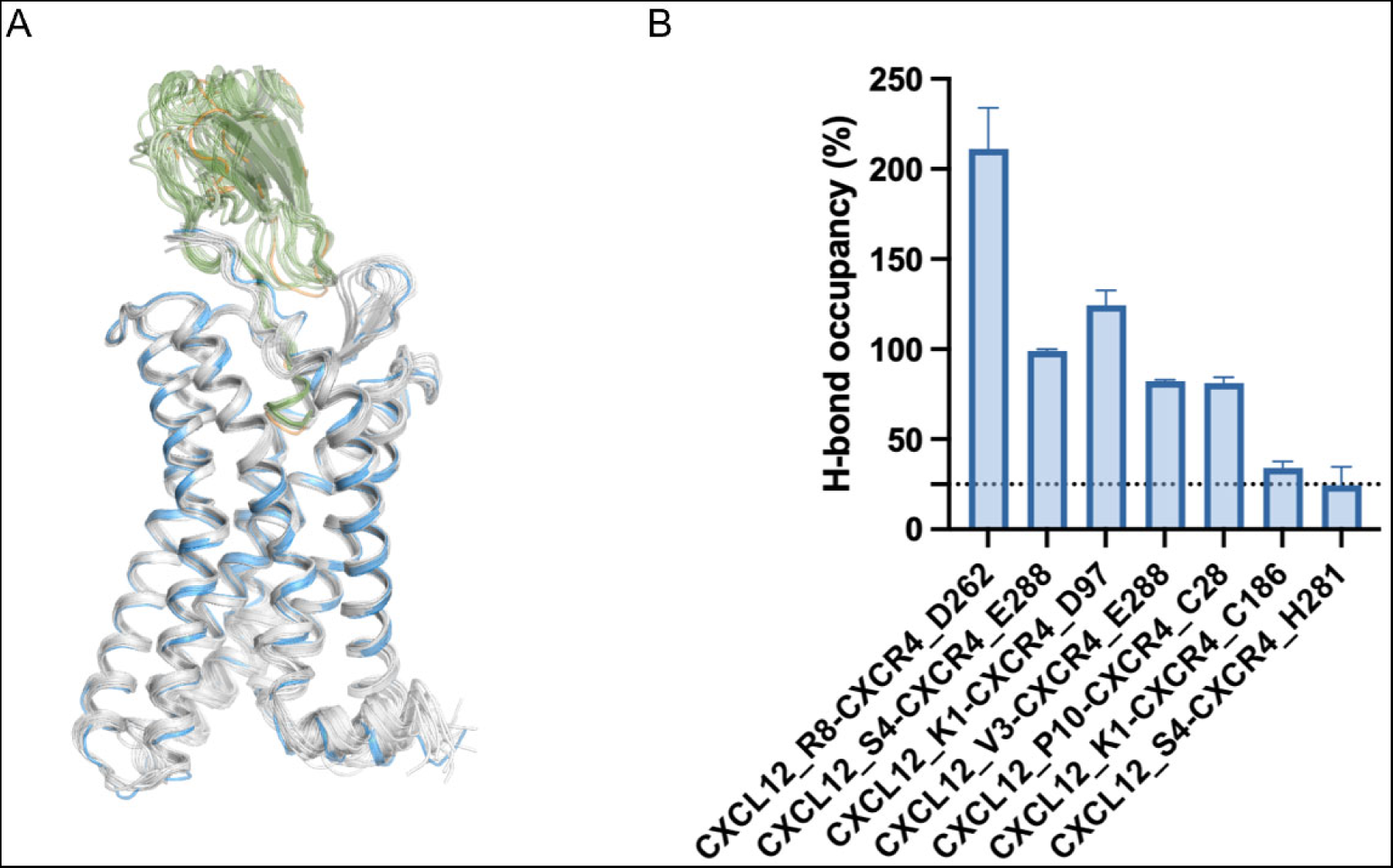
Molecular dynamics simulations of CXCL12-CXCR4 complex. **A** Representative snapshots of the complex collected during MD simulation superimposed with the cryo-EM model of CXCR4-CXCL12. **B** Hydrogen bond occupancy between CXCL12 and CXCR4 observed in MD simulations.

Hydrogen bond occupancy was analyzed at key binding residues (Fig. 4B). R8 of CXCL12 showed over 200% hydrogen bond occupancy with D262^6.58^ of CXCR4, outlining its importance in chemokine binding through salt bridge formation. The hydrogen bond occupancies among residue pairs of (CXCL12-CXCR4) S4-E288^7.39^, V3-E288^7.39^, and P10-C28^NT^ are greater than 50%, whereas the residue pairs K1-C186^ECL2^ and S4-H281^7.32^ bear a slightly lower hydrogen bond occupancy. Interestingly, the residue pair K1-D97^2.63^ show a relatively high occupancy of hydrogen bond, which is consistent with the findings from previous studies demonstrating the importance of D97^2.63^ in chemokine binding (Qin et al., 2015; Stephens et al., 2020; Wescott et al., 2016; Ziarek et al., 2017). However, the distance between D97^2.63^ of CXCR4 and K1 of CXCL12 is measured over 4 Å in our cryo-EM structure. This could be due to the flexibility of the lysine sidechain leading to the formation of salt bridge captured by MD simulation in the binding pocket of CXCR4. As the most important residue in CXCL12 for receptor activation, polar interactions of K1 with acidic residues C186^ECL2^ and D187^ECL2^ contribute more to chemokine ligand binding and receptor activation (Kufareva et al., 2017; Ngo et al., 2020; Stephens et al., 2020; Ziarek et al., 2017). Altogether, thermodynamic features of the CXCR4-CXCL12 complex as analyzed through MD simulations support our structural model.

### Activation mechanisms for the CXCL12-CXCR4-Gi complex

Next, we sought to identify details of CXCR4 activation from the perspective of conformational changes. CXCR4 shares a high sequence homology with another CXC chemokine receptor CXCR2, of which both the active and inactive state structures are available (Fig. S5). By comparing the active state structures of chemokine-bound CXCR2 (PDB ID: 6LFO) (Liu et al., 2020) and CXCR4 (this work), we found that CXCL12 mainly occupies the minor sub-pocket surrounded by TM1 and TM2 of CXCR4, while CXCL8 binds primarily to the major sub-pocket defined by TM5 and TM6 of CXCR2 (Fig. 5A). Moreover, CXCL12 takes a deeper insertion into the receptor binding pocket, as measured by the distance between the chemokine and the “toggle switch” W^6.48^ of the receptor. In fact, the 3.2 Å distance measured between V3 of CXCL12 and W^6.48^ of CXCR4 is among the shortest of all chemokine-chemokine receptor pairs in CRS2 (Urvas & Kellenberger, 2023). An inward movement of TM1 and an outward shift of ECL2 were also observed for CXCR4, accommodating a deeper minor sub-pocket for chemokine binding (Fig. 5B).

**Figure 5.**
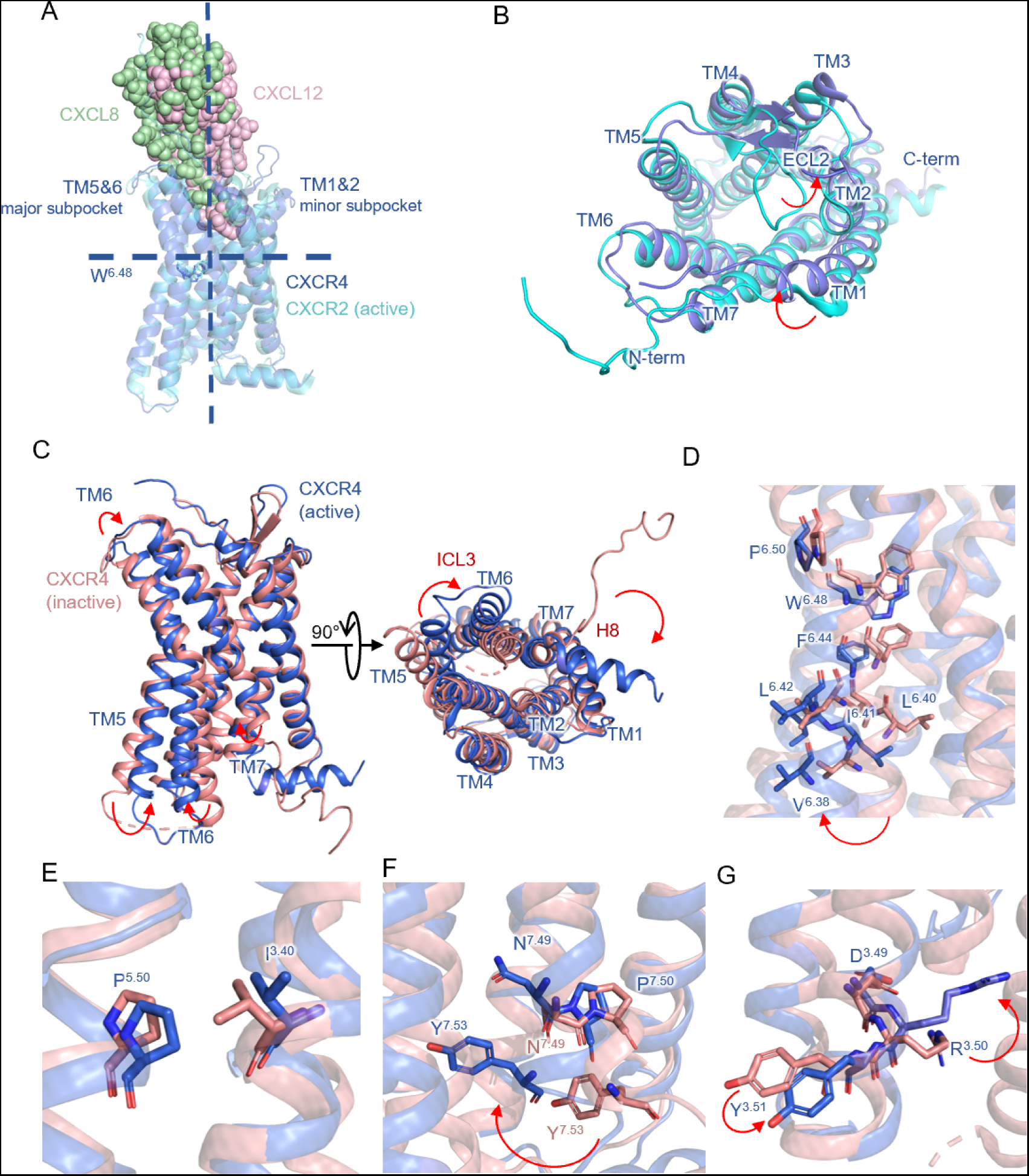
Activation of CXCR4-CXCL12 complex. **A** Comparison between the structures of CXCR4-CXCL12 and CXCR2-CXCL8. W^6.48^ is displayed as a reference of the insertion depth of chemokine ligands. Major and minor subpockets are separated by residues at position 3.32 and 7.39. **B** Extracellular view of superimposed structures of active CXCR4 (dark blue) and CXCR2 (cyan). **C** Comparison between the active CXCR4 (dark blue) and inactive CXCR4 (salmon) from side view (left) and intracellular view (right). **D** Comparison of TM6 residues between the active CXCR4 (dark blue) and inactive CXCR4 (salmon). **E** Comparison of P^5.50^-I^3.40^ motif between the active CXCR4 (dark blue) and inactive CXCR4 (salmon). **F** Comparison of NPxxY motif between the active CXCR4 (dark blue) and inactive CXCR4 (salmon). **G** Comparison of DRY motif between the active CXCR4 (dark blue) and inactive CXCR4 (salmon).

We further compared the differences between the inactive (PDB ID: 3ODU) (Wu et al., 2010) and active structures of CXCR4. In the active conformation, TM5 moves inwards with an outward displacement of the lower half of TM6 facing the intracellular compartment (Fig. 5C, left panel), supporting the interaction between the receptor and G protein heterotrimeric complex. The upper half of TM6 interfering with the extracellular environment shows an inward movement (Fig. 5C, right panel). This displacement may be particularly important for chemokine binding, as characterized by polar and nonpolar contacts between CXCL12 and CXCR4. Moreover, TM6 may link the chemokine binding and receptor activation through its conformational changes. Indeed, as suggested in a study that combined mutational library usage and computational simulations, a string of TM6 residues form a hydrophobic bridge in an active state model and link nearly all GPCR signaling motifs (Wescott et al., 2016). This feature of hydrophobic bridge is common among chemokine receptors and important for signal propagation and transmission by active CXCR4. We next examined the residues lining TM6. From P254^6.50^ to V242^6.38^, the distances between active and inactive structures increase from 2.0 Å to 5.0 Å (Fig. 5D). This outward swinging of TM6 in the active state of CXCR4 provides key space in order to accommodate the ⍺5 helix of G⍺i protein.

Besides, we specified the structural motifs for receptor activation. P^5.50^-I^3.40^ did not present a large movement (Fig. 5E). In comparison, the NPxxY motif, with a displacement of the intracellular end of TM7 to the center of G protein binding cavity of the receptor (Fig. 4C, left panel), demonstrates a significant movement inward (Fig. 5F). In transmembrane helix TM3, the DRY motif shows a counter-clockwise rotation, making R^3.50^ more stretched to the caveat for G protein binding. Therefore, the findings support a role for this particular motif in G protein association and transition of CXCR4 to an active state (Fig. 5G).

Next, to better characterize the interaction between an active state CXCR4 and G⍺i protein, we investigated the polar interactions in receptor-G protein interface. The ⍺5 helix of G⍺i protrudes into the receptor, surrounded by multiple polar contacts between D350 of G⍺i and K308^8.49^ of the receptor, C351 and R134^3.50^ of the receptor DRY motif, K353 and T240^6.36^, D341 and K234^6.30^, K314/E318 and K230^ICL3^ as well as E28 of G⍺i ⍺N helix and K149^3.38^ (Fig. 6A). Interestingly, TM6 of CXCR4 relays the conformational changes to the intracellular end, supporting a polar interaction network with Gi protein at T240^6.36^ and K234^6.30^, further stabilizing the receptor-G protein complex. Another pair of interaction between D350 of G⍺i and K308^8.49^ of the receptor Helix 8 stabilizes the orientation of Helix 8 in the active state (Fig. 6B). The outward displacement of ICL3 from inactive state to active state is supported by multiple contacts between K230^ICL3^ of CXCR4 and K314 and E318 of G⍺i (Fig. 6B). These conformational differences between active CXCR4 and inactive CXCR4 at receptor-Gi interface may contribute to the stabilization of CXCR4-Gi interaction. Ala substitution at R134^3.50^ and K149^4.38^ led to decreased G protein dissociation and cAMP inhibition (Fig. 6C and E). However, for other residues, especially residues lining TM6 for polar interaction network, the impact of introducing single point mutations is not sufficient to alter downstream G protein responses (Fig. 6D and F). These findings support our cryo-EM structure for an active state model of CXCR4 in complex with chemokine CXCL12 and Gi proteins.

**Figure 6.**
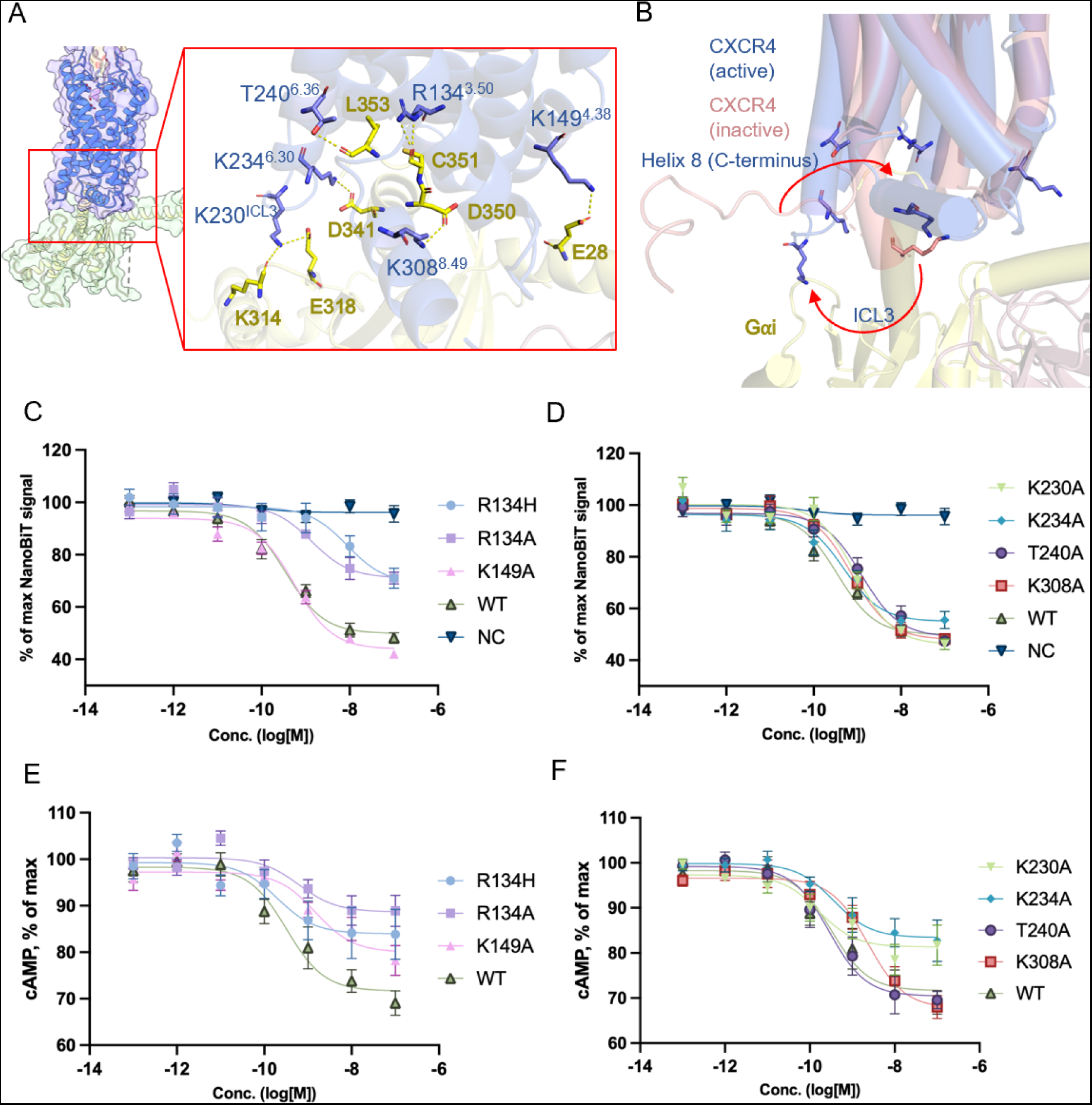
Interactions between CXCR4 and Gαi. **A** Polar interactions between CXCR4 (dark blue) and Gαi (yellow) highlighted in yellow dashes. **B** Comparison of active (dark blue) and inactive (salmon) CXCR4 structure at Gαi binding site. Red arrows indicate the conformational changes at Helix 8 and ICL3. **C-F** Mutations are introduced at Gαi-interacting residues of CXCR4. NanoBiT-based G protein dissociation assay (**C-D**) and cAMP inhibition assay (**E-F**) are performed for mutants. WT: wild-type CXCR4. NC: empty vector control. Data were collected from three independent experiments, each with three replicates.

### Comparisons of chemokine-chemokine receptor interactions at CRS1.5 and CRS2

Given the importance of CXCL12-CXCR4 interaction, we further compared CXCR4-CXCL12, CXCR2-CXCL8 and ACKR3-CXCL12 interaction at CRS1.5, CRS2 and position 7.39 of the receptor (Fig. 7). Common among these typical chemokine receptors, the classical “two-site” model applies to the interactions between chemokines and chemokine receptors. The N terminus of chemokine receptor forms CRS1 by binding to the chemokine N loop and β_3_ strand/40s-loop. CRS1 thus wraps around the globular core of the chemokine and is important for chemokine recruitment and stabilization (Szpakowska et al., 2012). Owing to its structural instability, CRS1 interaction is not visible in many chemokine-chemokine receptor structures (Urvas & Kellenberger, 2023). Several mutational studies identified that the negatively-charged N-termini of chemokine receptors bind to a shallow positively-charged crevice defined by the N-loop and β_3_ strand/40s-loop of the chemokine (Ishimoto et al., 2023; Shao et al., 2022; Stephens et al., 2020; Szpakowska et al., 2012; Yen et al., 2022; Zhang et al., 2021; Ziarek et al., 2017). This allows the distal end of N-terminus of chemokine receptors to stably wrap around the core of the chemokine. In a “two-step” model, CRS1 initializes the recognition and recruitment of chemokines. By dragging the chemokine ligand closer with its N-terminus distal end, CRS1 further stabilizes the chemokine-chemokine receptor complex and assists in the insertion of the chemokine to CRS2. Despite the lack of structural information at CRS1 in CXCR4-CXCL12 structure, many studies combining mutagenesis experiments and computational work have shed light on the mode of interaction at CRS1, as well as the role of CRS1 in chemokine recruitment and binding.

**Figure 7.**
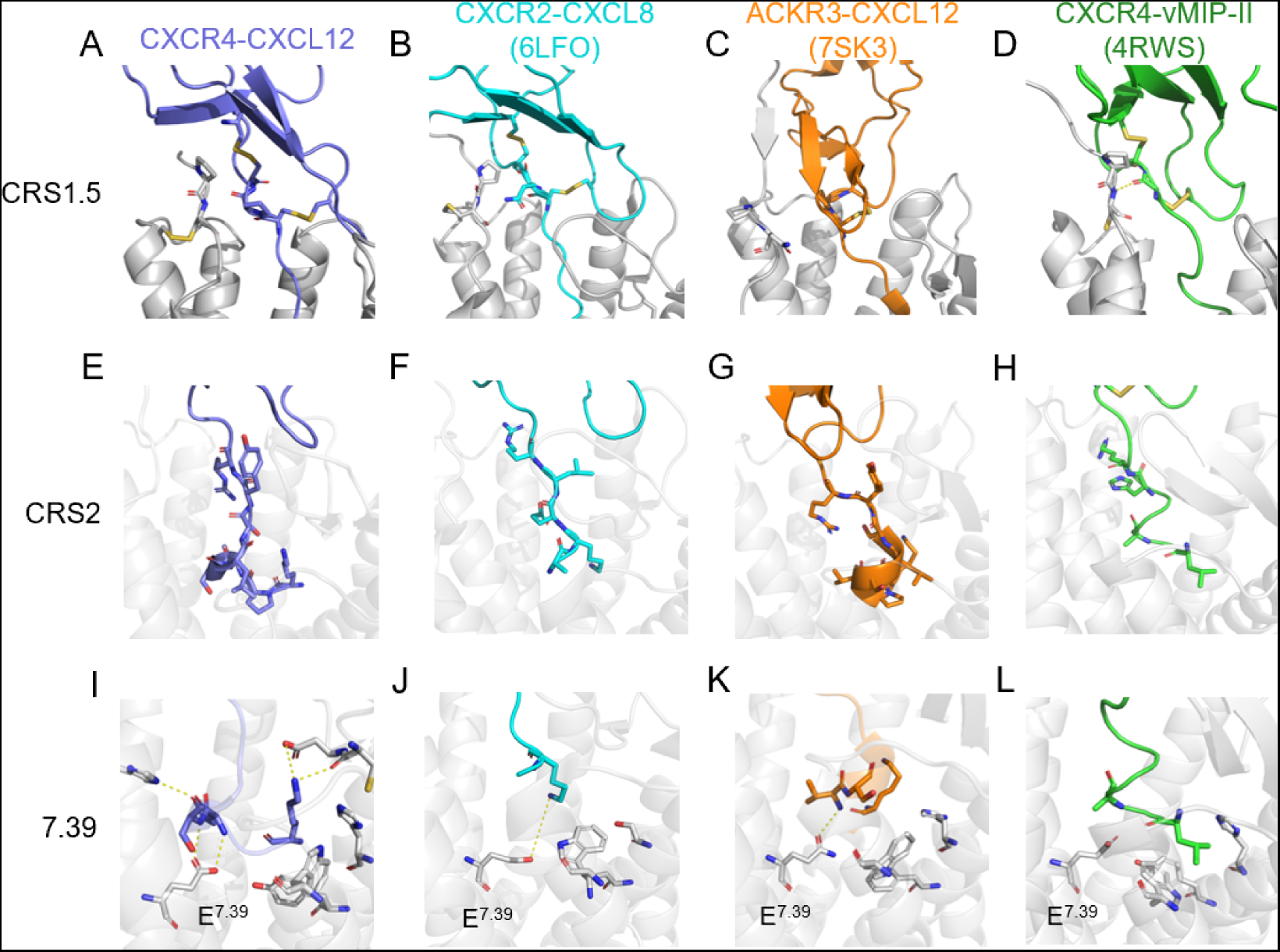
Comparisons of chemokine-chemokine receptor interaction at CRS1.5 and CRS2. **A-D** Interactions between receptor and chemokine at CRS1.5 are shown as sticks. **E-H** Interactions between receptor and chemokine at CRS2. Chemokine residues interacting with the receptor are displayed as sticks. **I-L** Interactions between receptor and chemokine at position 7.39 of the receptor. Interactions between position 7.39 and chemokine are highlighted in yellow dashes.

CRS1.5 is composed of a PC motif in the receptor N-terminus, closely interacting with the CXC motif conserved in CXC chemokines. This site of chemokine recognition was first proposed in the CXCR4-vCCL2 structure (Qin et al., 2015). The CRS1.5 interaction between PC motif and CXC motif is further enhanced by multiple disulfide bridges. The cysteine of the receptor N-terminus PC motif forms a disulfide bridge between N-terminus and TM7. The two cysteines in the CXC motif form two disulfide bridges linking CXC motif with 30s-loop and β3-strand of the chemokine, respectively (Fig. 7 A-D). This enhanced interaction at CRS1.5 is expected to orient the N-terminus of chemokine toward the TM cavity constituting CRS2, and the receptor N-terminus to CRS1, as a distinct feature of CXCRs but not present in CCRs (Shao et al., 2022; Urvas & Kellenberger, 2023; Wedemeyer et al., 2020; Zhang et al., 2021). However, CRS1.5 interaction is absent in atypical chemokine receptor 3 (ACKR3), as a scavenger receptor of chemokines (Fig. 7C) (Yen et al., 2022). As the core of CXCL12 turned around 80° when bound to ACKR3, the PC motif at chemokine N-loop goes closer to TM5 of ACKR3. At the typical site of CRS1.5 at the receptor N-terminus, the 30s-loop of CXCL12 forms few interactions with ACKR3. This may affect the receptor-chemokine interaction and explain the biased signaling property of ACKR3 when bound to CXCL12.

CRS2 is the canonical ligand binding pocket of chemokine receptors, where the N-terminus of chemokine inserts to activate the receptor. With a relatively large pocket, CRS2 can be divided into a major subpocket defined by TM3-7 and a minor subpocket formed by TM1, 2, 3, and 7 (Urvas & Kellenberger, 2023). Moreover, the depths of chemokine N-terminus insertion vary among CXCRs. In our model of CXCR4-CXCL12 interaction, the N-terminus of CXCL12 reaches the bottom of CRS2 binding pocket with multiple contacts including W^2.60^, Y^3.32^, Y^6.51^ and E^7.39^ (Fig. 7 E and I). This insertion is deepest among structures in our comparison. The N-terminus of CXCL12 has a turn at residue S4, allowing the first four amino acid residues to adopt a horizontal pose lying in the minor subpocket of CXCR4. This pose may maximize the interactions between CXCL12 and the bottom of CRS2, supporting its deep binding. CXCL8 activates CXCR2 with a very shallow insertion, yet its K3 side chain has ionic interaction with E^7.39^ (Fig. 7 F and J) (Liu et al., 2020). This shallow binding prefers multiple interactions with the major subpocket of CXCR2. For ACKR3-CXCL12 interaction, the N-terminus of CXCL12 inserts deep into CRS2 binding pocket and V3 has polar contact with E^7.39^ (Fig.7 G and K) (Yen et al., 2022). While the N-terminal residues of CXCL12 mainly occupy the minor subpocket, K1 of CXCL12 reaches out into the major subpocket of ACKR3. This difference may correlate with the ∼80° turn of CXCL12 globular core when bound to ACKR3. CXCR4 binds to a viral chemokine antagonist vCCL2 (Qin et al., 2015). The binding between CXCR4 and vCCL2 is shallow compared to CXCR4-CXCL12 binding (Fig. 7 H and L). Similar with CXCL12, vCCL2 occupies the space in the minor subpocket of CXCR4, and no interaction presents between vCCL2 and E^7.39^ of CXCR4. Given the importance of CRS2 interactions in receptor activation and downstream signaling, understanding the roles of chemokine N-terminal residues in CRS2 interaction may be crucial for future development of CXCR4 agonists and antagonists.

In summary, the present work presents a high-resolution structure of CXCR4 bound to its native agonist, CXCL12, in monomeric form. The availability of this structure allows comparison with (and confirmation of) the predicted CXCR4 structures as well as its interactions with the orthosteric and viral chemokines. These previous studies employed NMR, molecular modeling, disulfide crosslinking and systemic mutagenesis approaches, generating a wealth of insights that have helped our understanding of this important chemokine-chemokine receptor interaction. With the structural information of CXCL12-CXCR4-Gi complex, additional ligands may be designed for further manipulation of myeloid cell trafficking and homeostasis.

## Acknowledgments

This work was supported in part by grants from the Science, Technology and Innovation Commission of Shenzhen Municipality GXWD20201231105722002-20200831175432002 (R.D.Y. and L.Z.), JCYJ20200109150003938 (L.Z.), RCYX20200714114645019 (L.Z.), and Shenzhen Science and Technology Program Grant No. RCBS20221008093330067 (A.L.). This work was also supported by National Natural Science Foundation of China 31971179 (L.Z.), and 32070950 (R.D.Y.), China Postdoctoral Science Foundation 2022M713049 (A.L.), the Ganghong Young Scholar Development Fund (R.D.Y.), the fund from Kobilka Institute of Innovative Drug Discovery at The Chinese University of Hong Kong, Shenzhen (R.D.Y.), and the fund from Shenzhen-Hong Kong Cooperation Zone for Technology and Innovation (HZQB-KCZYB-2020056). The authors thank the Kobilka Cryo-Electron Microscopy Center in The Chinese University of Hong Kong, Shenzhen for cryo-electron microscopy analysis, and Warshel Institute for Computational Biology (funds from Shenzhen City and Longgang District) for computational work.

## Author Contributions

R.D.Y., Y.L., and A.L. conceived, designed and initiated the project; A.L. cloned the constructs, expressed and purified the protein complex, and determined the cryo-EM structure; Y.L. conducted mutagenesis analysis, functional assays and structural comparison; A.L., Y.L., X.L., Q.L., and L.Z. performed MD simulations; Y.L. and W.Z. conducted flow cytometry experiments and signaling assays; L.Z. and R.D.Y. supervised the research; Y.L., A.L., and R.D.Y. analyzed data and wrote the manuscript with input from all authors.

## SUPPLEMENTARY INFORMATION

**Fig. S1.**
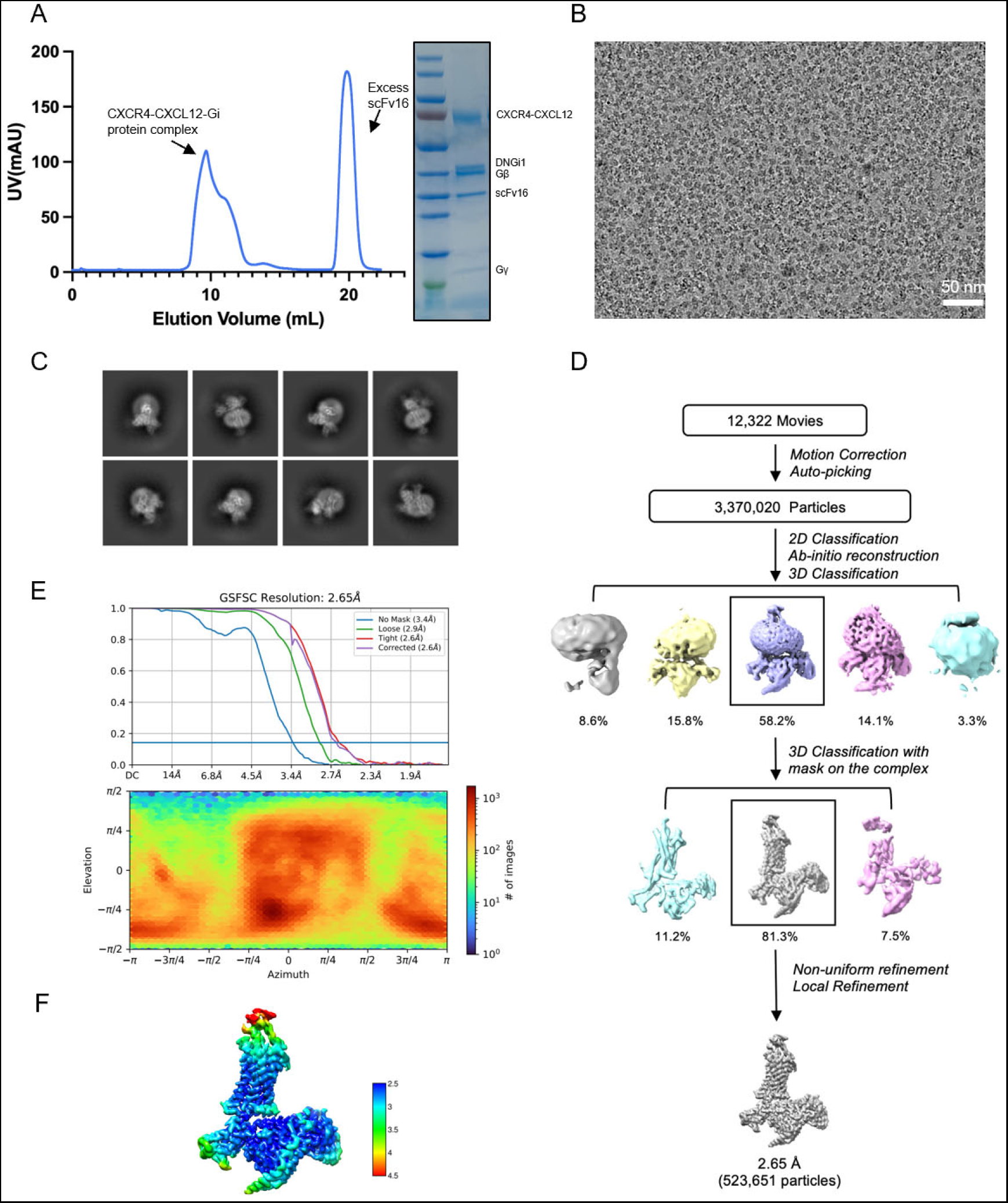
Protein purification, cryo-EM data collection, structure determination and cryo-EM maps. **(A)** Size-exclusion chromatography elution profiles of the C9-GPR1-G protein complex and the SDS–PAGE and Coomassie blue staining of the C9-GPR1-G protein complex. **(B)** Representative micrograph of the complex particles from 3,609 movies. **(C)** Representative 2D averages. **(D)** Workflow for cryo-EM image processing. **(E)** Gold standard Fourier shell correlation (FSC) curve indicates overall nominal resolution at 2.65 Å using the FSC=0.143 criterion. **(F)** Local resolution map.

**Fig. S2.**
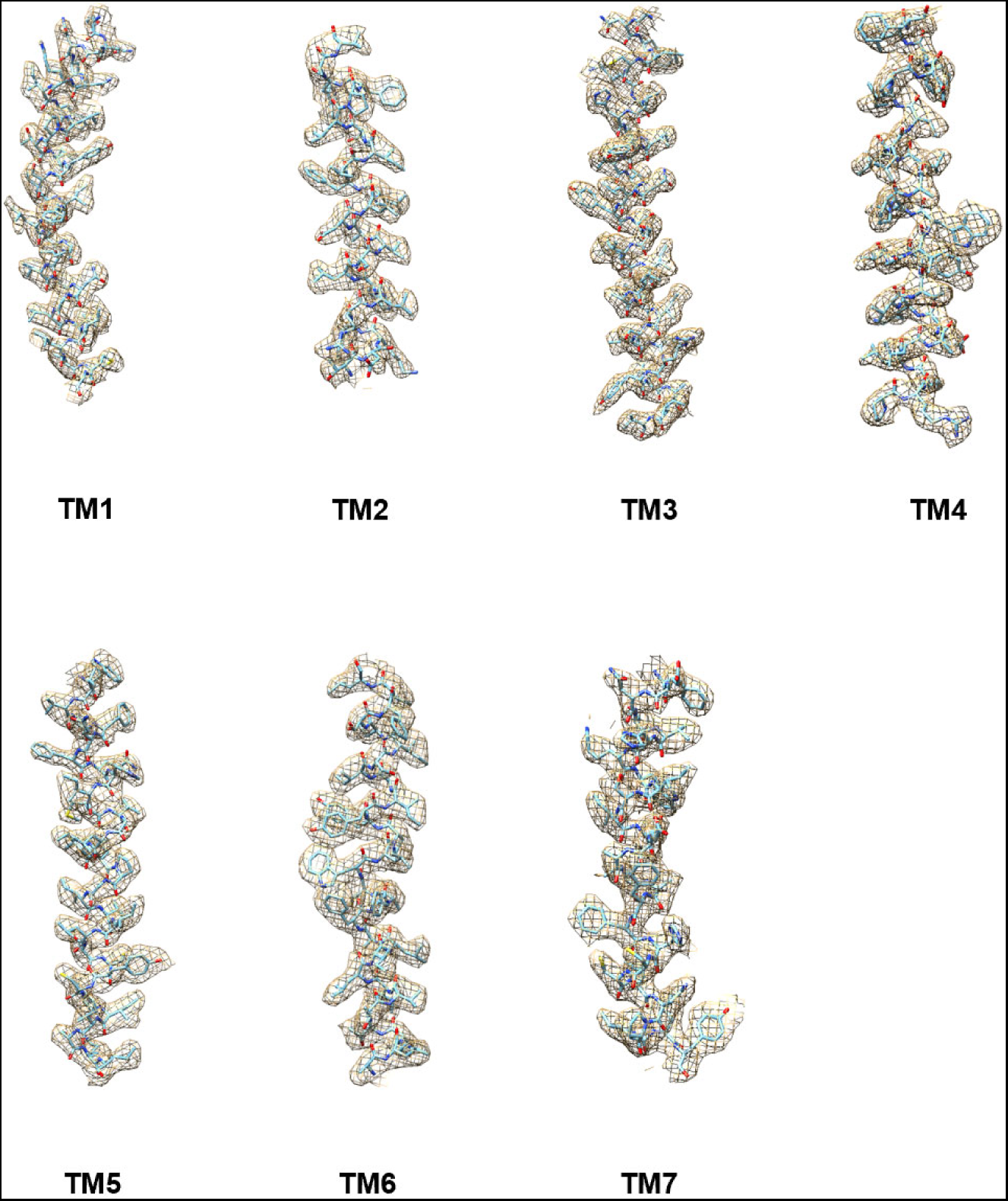
Representative density maps and models for TM1-7.

**Figure S3.**
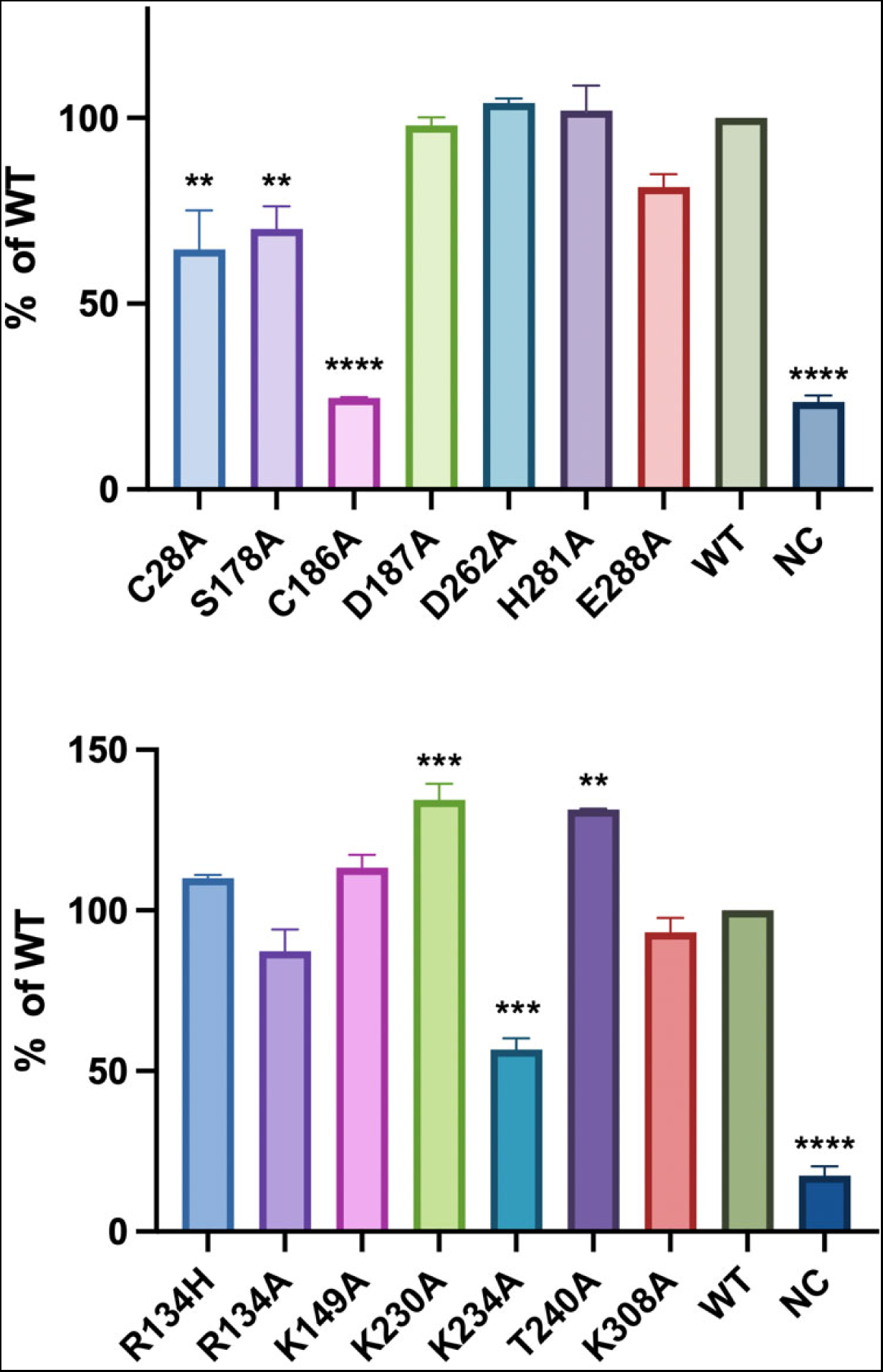
Cell surface expression of CXCR4 mutants. HEK293T cells were transfected WT or mutant CXCR4 for 24 h at 37°C. Cells were then incubated with an APC-conjugated CXCR4 antibody (12G5) for 30 mins on ice. The fluorescence signals on the cell surface were quantified by flow cytometry. Data shown are means ± SEM from 3 independent experiments. *, *p* < 0.05.

**Figure S4.**
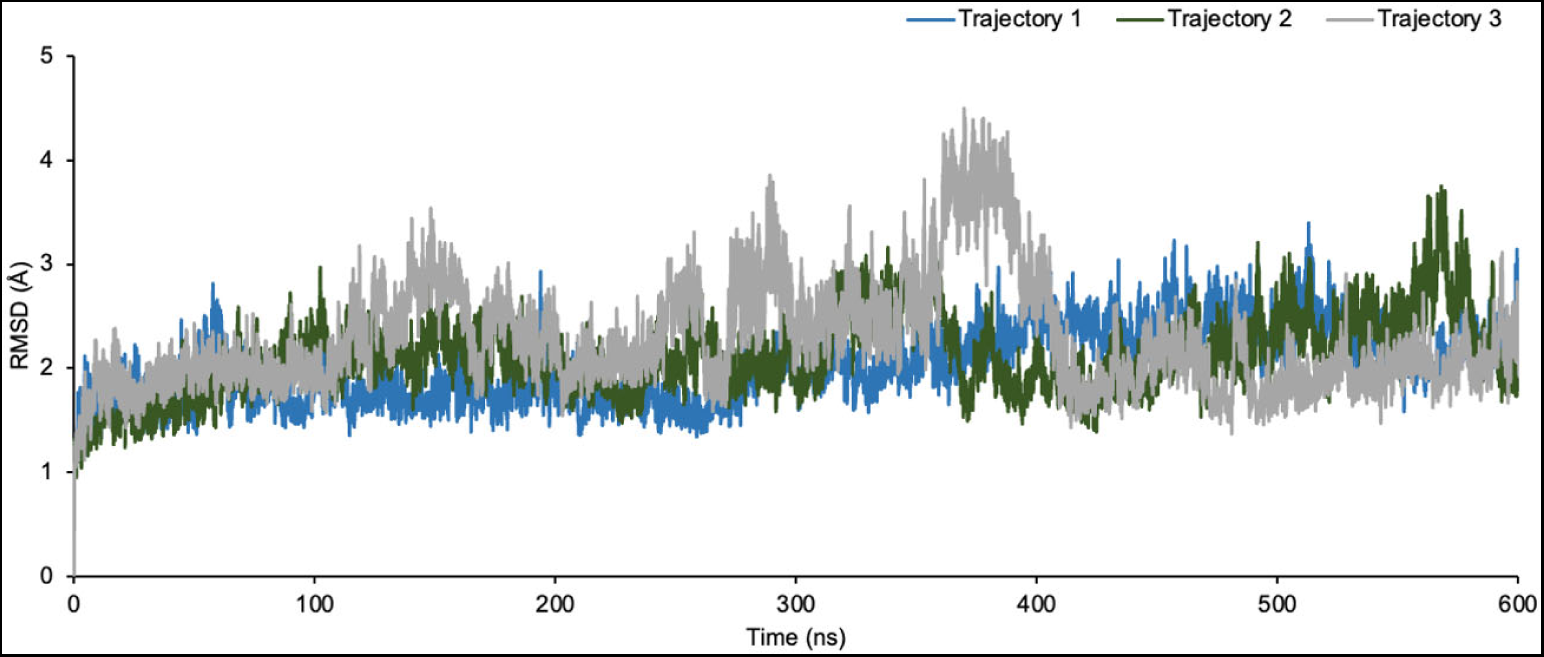
Root mean square deviations (RMSD) of the backbones on the receptor and the chemokine ligand throughout each trajectory from three independent 600 ns MD simulation.

**Figure S5.**
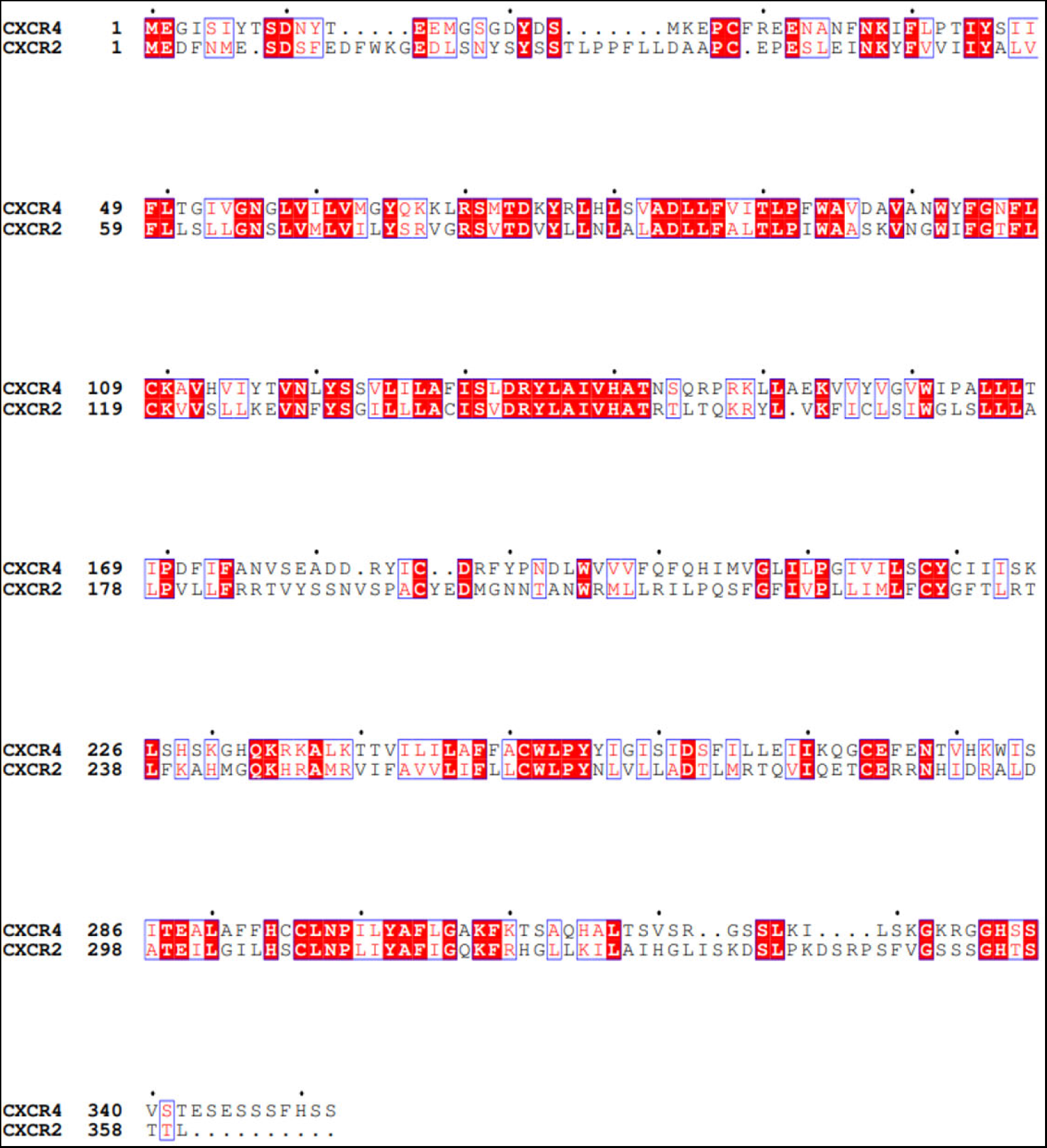
Sequence alignment between CXCR4 and CXCR2.

**Table S1.** Cryo-EM data collection, model refinement and validation statistics.

| **Data collection and processing** | **CXCL12-GXCR4-G<sub>i</sub>-scFv16** |
| --- | --- |
| Magnification | 105000 |
| Voltage (kV) | 300 |
| Electron exposure (e-/Å<sup>2</sup>) | 52.0 |
| Defocus range (μm) | -1.2~-2.5 |
| Pixel size (Å) | 0.85 |
| Symmetry imposed | C1 |
| Initial particle projections (no.) | 3,370,020 |
| Final particle projections (no.) | 523,651 |
| **Refinement** |  |
| Initial model used (AlphaFold code) | AF-P61073-F1 |
| Map resolution (Å) | 2.65 |
| FSC threshold | 0.143 |
| **Model composition** |  |
| Non-hydrogen atoms | 9536 |
| Protein residues | 1217 |
| **R.m.s. deviations** |  |
| Bond lengths (Å) | 0.003 |
| Bond angles (°) | 0.624 |
| **Validation** |  |
| MolProbity score | 2.12 |
| Clashes core | 7.54 |
| **Ramachandran plot** |  |
| Favored (%) | 95.75 |
| Allowed (%) | 4.25 |
| Outliers (%) | 0.00 |

